# Pervasive phenotypic impact of a large non-recombining introgressed region in yeast

**DOI:** 10.1101/2020.01.29.925289

**Authors:** Christian Brion, Claudia Caradec, David Pflieger, Anne Friedrich, Joseph Schacherer

**Author notes:** Correspondence to: JS.

## Abstract

To explore the origin of the diversity observed in natural populations, many studies have investigated the relationship between genotype and phenotype. In yeast species, especially in *Saccharomyces cerevisiae*, these studies are mainly conducted using recombinant offspring derived from two genetically diverse isolates, allowing to define the phenotypic effect of genetic variants. However, large genomic variants such as interspecies introgressions are usually overlooked even if they are known to modify the genotype-phenotype relationship. To have a better insight into the overall phenotypic impact of introgressions, we took advantage of the presence of a 1-Mb introgressed region, which lacks recombination and contains the mating-type determinant in the *Lachancea kluyveri* budding yeast. By performing linkage mapping analyses in this species, we identified a total of 89 loci affecting growth fitness in a large number of conditions and 2,187 loci affecting gene expression mostly grouped into two major hotspots, one being the introgressed region carrying the mating-type locus. Because of the absence of recombination, our results highlight the presence of a sexual dimorphism in a budding yeast for the first time. Overall, by describing the phenotype-genotype relationship in the *L. kluyveri* species, we expanded our knowledge on how genetic characteristics of large introgression events can affect the phenotypic landscape.

## Introduction

Phenotypic diversity, such as variation of fitness, sensitivity to diseases, or differences of behavior, is intrinsically linked to genetic variation. Consequently, in health research, food industry and many other biological field, the genotype-phenotype relationship is deeply investigated to have a better understanding of the cause of this diversity and its evolution. Such relationship can be dissected by using large genotyped and phenotyped populations (1000 Genomes Project Consortium et al. 2010; UK10K Consortium et al. 2015; Alonso-Blanco et al. 2016; Peter et al. 2018). In yeast, and especially in the model species *Saccharomyces cerevisiae*, these populations are generally constituted of recombinant offspring originated from crosses, which prevent biases of the association analysis regarding population structure, or the effect of rare variants (Brem et al. 2002; Steinmetz et al. 2002; Fay 2013; Bloom et al. 2019; Fournier et al. 2019). These analyses allowed the identification of a large number of quantitative trait loci (QTL), responsible for the variation of a broad set of phenotypes such as the growth fitness in various conditions, cell morphology, or gene expression (RNA level, eQTL, and protein level, pQTL) (Brem et al. 2002; Nogami et al. 2007; Smith and Kruglyak 2008; Bloom et al. 2013; Fay 2013; Albert et al. 2014; Clément-Ziza et al. 2014; Albert et al. 2018; Peltier et al. 2019).

Most of these analyses mainly focused on identifying the effects of punctual mutations. However, other types of genetic variants have been shown to affect the phenotypic diversity and the dynamics of species evolution. One of these variants corresponds to the transfer of genetic material across species, such as introgression events. Such transfers have been identified in yeast and can confer phenotypic advantages in wine making process (Novo et al. 2009; Marsit et al. 2015). In addition, it has been shown through the lens of the sequences of 1,011 *Saccharomyces cerevisiae* genomes that these introgression events are common. In total, 885 introgressed genes coming from the *Saccharomyces paradoxus* sister species have been identified in this population (Peter et al. 2018). However, the overall impact of introgression events on the dynamic of evolution of yeast species remain to be explored.

While most of the introgression events are usually small in *S. cerevisiae*, the pre-duplicated species *Lachancea kluyveri* carry a very large (∼ 1 Mb) genomic introgression, corresponding to the left arm of the chromosome C (Génolevures Consortium et al. 2009; Payen et al. 2009). This introgressed region is common to all the sequenced isolates of the species (Friedrich et al. 2015). It presents a higher GC content compare to the rest of the genome (53% *vs*. 40%) and displays different evolutionary features: the genetic diversity is higher (π = 0.019 *vs*. π = 0.017), the linkage disequilibrium is shorter (LD_1/2_ = 0.3 kb *vs*. LD_1/2_ = 1.5 kb), and the mutation types are unbalanced (Friedrich et al. 2015). Recently a donor species, *Eremothecium gossypii*, has been proposed as the donor species, based on synteny and codon usage (Landerer et al. 2019). We also observed that no double strand breaks occurred in this region during meiosis, preventing cross-overs and allelic shuffling (Brion et al. 2017). This latest feature, associated to the fact that the region contains the mating type locus (*MAT* locus), is particularly noteworthy. Indeed, the region around the *MAT* locus is generally associated with a diminution of the recombination rate in yeast, but it is usually restricted to a very small portion of the genome around that locus.

To have a global view of the genetic architecture of traits in *L. kluyveri* as well as to assess the phenotypic impact of this large non-recombining introgressed region, we performed linkage mapping on large populations of segregants. Quantitative growth in 64 conditions, which induce various physiological and cellular responses, and gene expression variations were measured, leading to the determination of 89 QTL and 2,187 eQTL, mostly grouped into two common major hotspots. Interestingly, one of the QTL and eQTL hotspots corresponds to the introgressed region, highlighting the pervasive phenotypic impact of it. We also demonstrated an association between different traits and the mating-type locus due to the absence of recombination in the introgressed region. This observation implies a sexual differentiation in the species and consequently highlights the presence of a sexual dimorphism in a budding yeast for the first time. Finally, by providing an exhaustive view of the QTL and eQTL landscape, we were able to compare the genotype-phenotype maps across different yeast species.

## Results

### *Lachancea kluyveri* introgressed region shows no recombination after two cycles of meiosis

In order to identify the genetic basis underlying the phenotypic diversity in *L. kluyveri*, we used a cross between the NBRC10955 (*MAT*a), and 67-588 (*MAT*α) strains, for which the density of punctual genetic variant is around 7 mutations per kb (0.7% of diversity). In our previous survey focusing on the recombination landscape of this species, we generated and sequenced 196 F1 offspring (from 45 full tetrads) (Brion et al. 2017). Here, we used 180 of these strains, plus two additional sequenced strains that do not come from a full tetrad. We genotyped this F1 population by defining the parental origins of 56,612 variants along the genome and identified the position as well as the number of recombination events per spore. These recombination events are the result of cross-overs during meiosis but also of loss of heterozygosity, likely induced by incomplete meiosis and return to growth event prior the final meiosis (Brion et al. 2017). On average, 14.7 recombination events were detected by strains, ranging from 3 to 30 (figure S1), while none were observed in the introgressed region located on the left arm of the chromosome C.

Because the recombination rate in *L. kluyveri* is 3.7 time lower than in *S. cerevisiae* (Brion et al. 2017), we generated a F2 population of recombinant strains from the same cross in order to perform an eQTL analysis. We crossed *MAT*a and *MAT*α strains from the F1 population (69 crosses). These hybrids were put on sporulation condition and the resulting tetrads were dissected. The F2 populations is composed of 50 segregants coming from 50 independent crosses and full tetrads (one spore per tetrad was selected). RNA sequencing (RNA-seq) of this F2 population allowed characterization of the parental allele inheritance for each segregant. By only using variants in coding frame, the number of used polymorphic sites was reduced to a total of 37,529. Because recombination is low, we used pseudo-markers every 3 kb to reduce the polymorphic sites to 3,779, which still allowed us to detect recombination accurately. Again, in this population, no recombination events were detected in the introgressed region, inducing perfect linkage across its variants. The average number of recombination events per spore in the F2 population was 19.5, ranging from 10 to 32 (figure S1). As expected, the number of recombination events was higher than in the F1 population, but not twice as much as undetected recombination events could occur in the homozygote regions of the F1 hybrids. Due to the lower recombination rate in *L. kluyveri* compared to *S. cerevisiae*, we observed larger blocks of linked variants along the genome (figure S2).

### A significant part of the heritable phenotypic variance is linked to the mating-type locus

To explore the genetic basis of phenotypic diversity in *L. kluyveri*, we looked at the variation of fitness and expression in our F1 and F2 populations, respectively. In this study, we defined fitness as the individual growth capacity in a specific condition. The fitness across 64 various conditions was estimated by measuring the colony size of the 182 F1 strains growing on solid media. These conditions include temperature variation, various carbon and nitrogen sources, the presence of toxic compounds and pH variation (table S1 and S2). From these data, we first estimated the phenotypic variance as well as the broad-sense heritability (*H*^*2*^) and we observed that most of the phenotype display a *H*^*2*^ superior to 70% (60 out of 64 tested conditions). This clearly indicates that the experimental error is low and most of the variation is due to heritable factors (figure 1A). We then looked at the distribution of the traits in the population and almost all the phenotypes displayed a normal distribution, suggesting a complex genetic control (examples in figure 1B-D). Only four phenotypes displayed a bimodal distribution, which could indicate a major locus control (6-azauracil 600 mg/L, anisomycine 50 mg/L, NaCl 1M and 1.5M) (figure 1C). We also quantified the mRNA for 5,380 annotated genes across the 50 strains of the F2 population using the RNA-seq data, and estimated the *H*^*2*^ for each gene using expression data from a previous experiment (see method). The expression of the majority of the genes (∼ 78%) showed a *H*^*2*^ greater than 70%, indicating that most of gene expression variation is carried by heritable factors.

**Figure 1.**
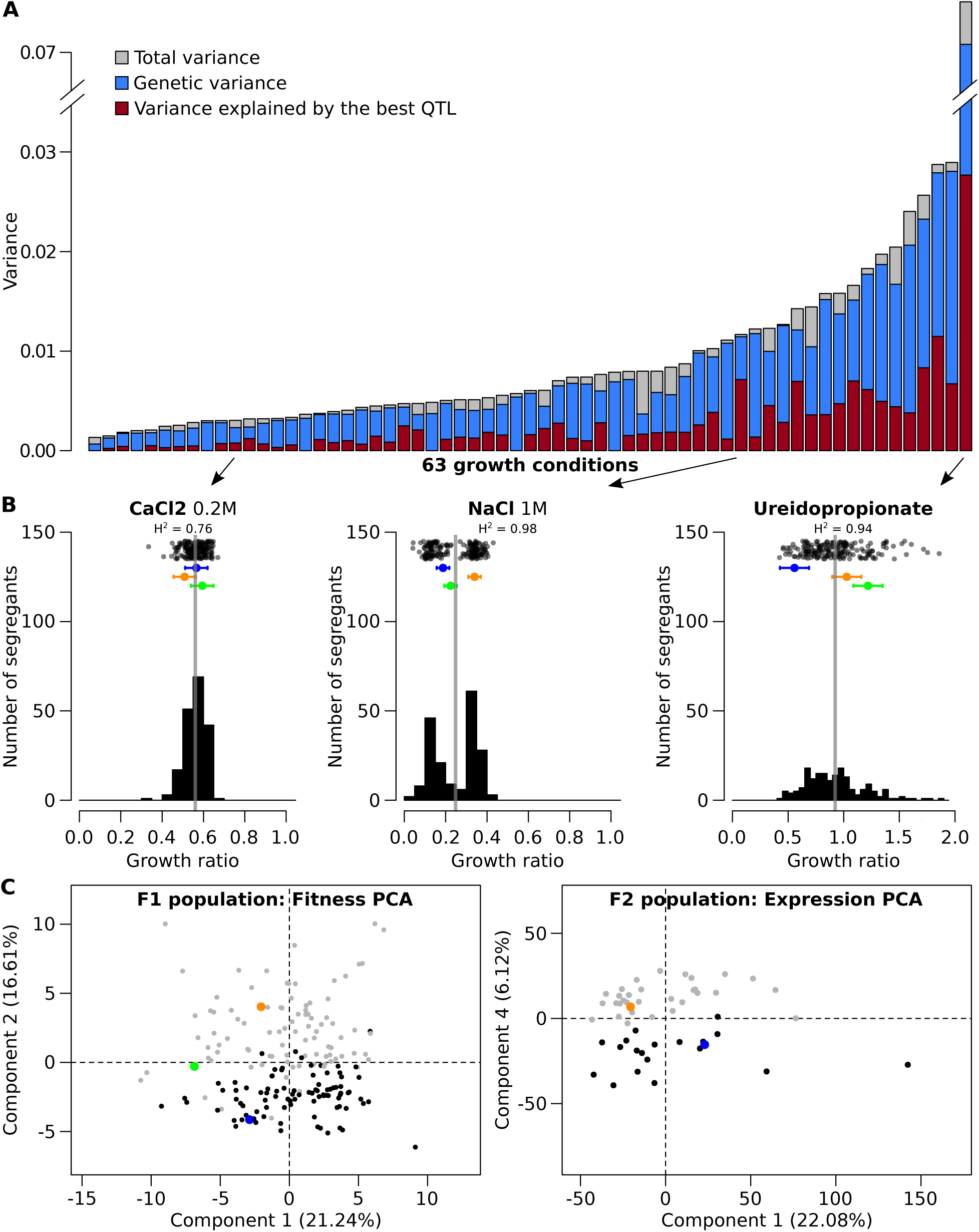
Phenotypic diversity in segregating populations. A) Distribution of the total variance of the growth ratio across the 63 conditions (YPD colonies sized are excluded), compared to the genetic variance and the variance explained by the best QTL. B) Three examples of distribution of the growth ratio in the F1 population selected from figure 1A (indicated by the arrows), for growth in 0.2M of CaCl_2_ (normal distribution), 1M of NaCl (bimodal distribution), and Ureidopropionate as unique nitrogen source (unclear distribution with a large variance). Top of each plot are the dot clouds distribution of the phenotype in the population, where mean value of the segregants are represented by grey lines. Means across the replicates of the parental strains are: NBRC10955 (blue), 67-588 (orange), and hybrid NBRC10955×67-588 (green). C) Principal component analysis of the fitness measure for F1 population (left) and expression profiles of the F2 population (right). Both populations are separated according to the *MAT* genotype: *MAT*a genotype (black) and *MAT*α genotype (grey). Parental strains follow the same color code as figure 1B.

We explored the phenotypic dispersion of the strains in both the F1 and the F2 population using principal component analyses (PCA) (figure 1E). The projection of the strains in the two PCA showed that our data have no outliers with extreme phenotypes and that the parental strains are generally separated, *i.e.* behaving differently. More importantly, we observed that for both PCA, a component can separate the populations accordingly to their mating-type: the second component (16.6%) for the fitness phenotypes in the F1 population, and the fourth component (6.12%) for gene expression in the F2 population (figure 1E). This result clearly highlighted a link between the *MAT* locus and a large part of the phenotypic variance. This link was also observed in a clustering analysis, regrouping genes based on their expression profiles across segregants. As expected, clustered genes were involved in similar biological processes due to the link between regulatory network and biological function (Figure S3). However, a cluster containing genes involved in the mating-type determination was also constituted of genes involved in other unrelated processes, and was enriched in genes located in the introgressed region. Overall, the variation of fitness and expression was partly associated to the mating type, which suggested an important role of the introgression carrying the *MAT* locus in *L. kluyveri* diversity. Such link was confirmed by the following QTL analysis.

### Majority of the detected QTL in *L. kluyveri* are localized within two pleiotropic loci

Using genotypic and phenotypic data of the F1 population, we identified loci that have an impact on fitness variance. The linkage analysis allowed detection of 89 QTL for 90% of the fitness measures (58 out of the 64 growth condition), with a false discovery rate (FDR) of 5%. The number of significant additive QTL detected per phenotype vary from one to seven (figure 2A and S4, table S3). Similarly, in the F2 population, we linked gene expression variation to the genotype and detected 2,187 eQTL for 2,048 genes (about 34% of the genome), at a logarithm of odd (LOD score) threshold of 3.7 (FDR of ∼ 4%, Figure S5). Only one eQTL was detected for most of the genes (1,911), while 135 genes have two eQTL and only two genes have three eQTL. Due to the low recombination rate in *L. kluyveri*, the loci identified are large, with a median of 70 kb, encompassing approximately 25 genes (figure 2B and S6, table S4).

**Figure 2.**
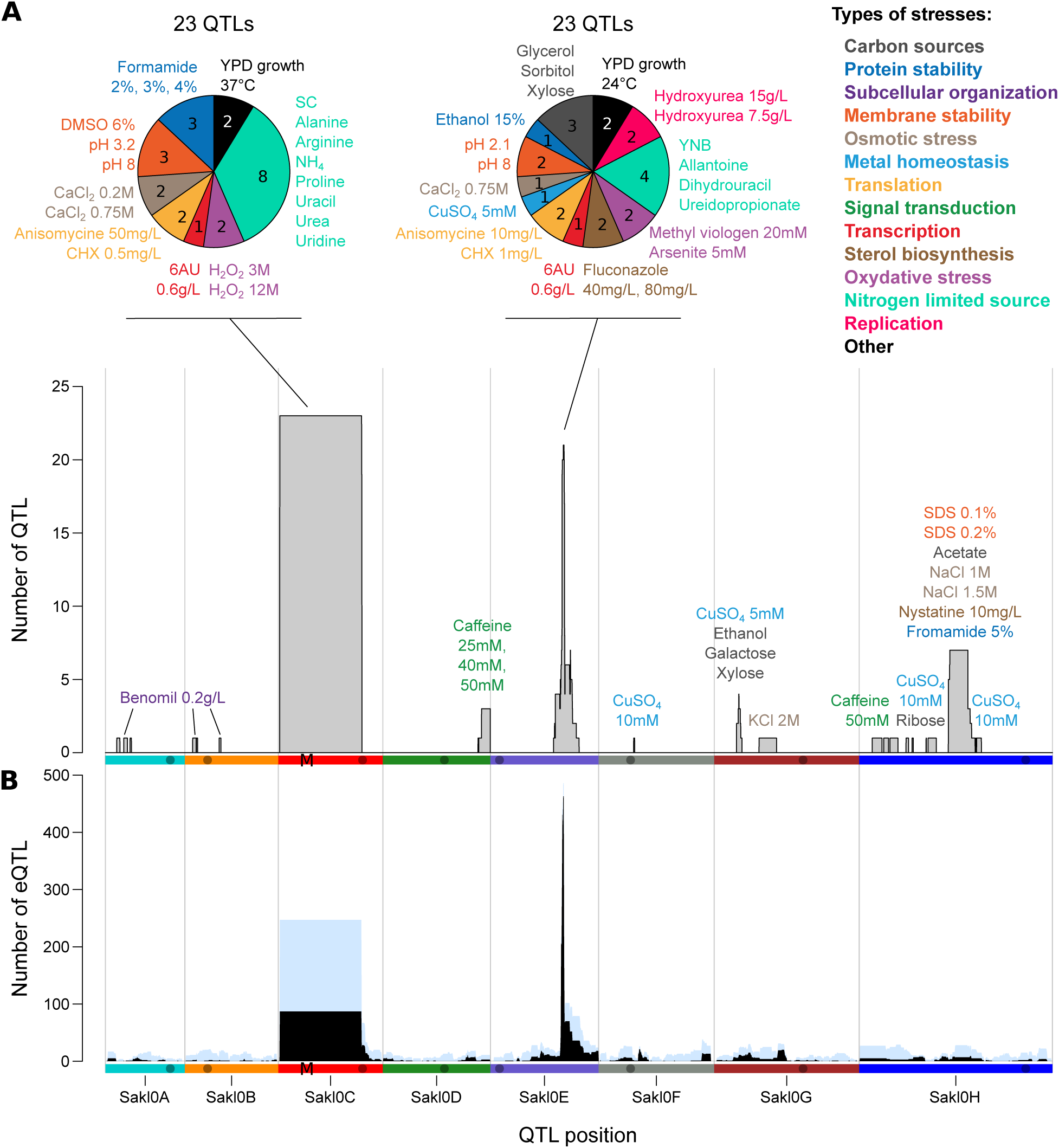
Summary of the linkage analysis. A) Density of fitness QTL (represented by grey bars) detected along the genome. The pie charts describe the two major QTL hotspots, indicating the number of growth conditions inducing different types of stresses. B) Density of distant eQTL (in black) and all eQTL (distant and local, in light blue) along the genome. The M indicates the position of the *MAT* locus.

We estimated the variance explained by each QTL and found that they explain a least 7.3% of the variance and only one case explain more that 60% (figure 1A and S7A). Similarly, we calculated the variance of gene expression explained by each eQTL and observed that these values can not be lower than 20%. This indicated a lower detection power for the eQTL compared to the fitness QTL, which can be explained by the smaller size of the F2 population. A major allelic effect was observed for 400 eQTL, explaining more than 60% of the expression variance (Figure S7B). The only strong genetic control on fitness detected affects resistance to sodium chloride (NaCl, 1M) and explains perfectly the bimodal distribution of the phenotype (61.4% of variance explained). This QTL is located on chromosome H and is due to a variation in the *SAKL0H13222g* gene, an ortholog of the *S. cerevisiae ENA2* gene. This gene encodes a membrane sodium transporter and has already been shown to have an impact on sodium resistance in our cross (Sigwalt et al. 2016). Interestingly, the *ENA2* allele also affects fitness in other media containing sodium (SDS, sodium acetate). An eQTL explaining 94% of the expression variance of *ENA2* is also located at this position, suggesting that the genetic variant acts through a modification of the expression of the *ENA2* gene. Similarly to *L. kluyveri*, the modification of *ENA2* expression, via copy number variation, also leads to the variation of salt tolerance across *S. cerevisiae* strains (Ruiz and Ariño 2007; Doniger et al. 2008).

We then examined the location of the 89 fitness QTL and 2,187 eQTL across the genome and found a large number of these QTL are regrouping in only two genetic locations, *i.e.* two hotspots (Figure 2). The first hotspot is located on the chromosome C, covering the introgressed region, and is composed of 23 fitness QTL and 247 eQTL. The second hotspot is on the chromosome E and contains 23 fitness QTL and 512 eQTL. The presence of QTL hotspots is a phenomenon observed in similar analyses in yeast, generally due to one genetic variant with pleiotropic effects, affecting multiple networks and traits. With the extreme detection power of more than 1,000 segregants, 9 fitness QTL and 102 eQTL hotspots were determined in *S. cerevisiae*, unveiling how frequent these pleiotropic variants are in this species (Bloom et al. 2013; Albert et al. 2018).

### The absence of recombination is responsible for the genetic link between the *MAT* locus and a large number of traits

The overlap between the fitness and expression QTL hotspots and the introgressed locus demonstrates an important role of this region on the phenotypic diversity. Because of the absence of recombination, all these QTL cover the large 1-Mb region, comprising of approximately 500 genes. Therefore, it is difficult to know if these QTL are due to one pleiotropic causal variant or multiple genetically linked variants. However, our results are in favor of the latter hypothesis, as the fitness QTL detected affect growth in very different types of stress, including membrane stability, oxidative stress, and nitrogen limitation. Similarly, the eQTL hotspot is probably due to multiple causal genetic variants, as they impact genes with unrelated functions. In addition, most of the eQTL detected in the introgressed region affects the expression of genes located within this region and are probably due to variants in *cis*-regulatory elements. By considering only genes located outside of this region, a weak enrichment in amino acid biosynthesis function (FunSpec p-value: 1.5e^-6^) was found for the fitness QTL affecting growth on several nitrogen limited media. Additionally, as expected, we found an enrichment for genes involved in peptide pheromone maturation (FunSpec p-value: 7.9e^-5^), explained by the presence of the mating-type regulator in the introgressed region.

Additionally, we observed another important consequence of *MAT* locus being present in the non-recombinant region. All the growth traits and gene expression impacted by variants in this region are genetically linked to the mating-type, which explains the results of the PCA analysis (figure 1E). It also means that these variants are hitchhiking the selection of the *MAT* alleles, necessary for mating. This adds up to a general consequence of absence of recombination: a possible accumulation of deleterious mutations in this region. This was confirmed using the average dN/dS values for each gene across the 28 sequenced *L. kluyveri* natural isolates (Friedrich et al. 2015). The dN/dS value corresponds to the ratio of density of non-synonymous polymorphism reported to the density of synonymous polymorphism. This value is generally higher for the genes located in the introgressed region, including for genes highly conserved across species (figure S8) (Brion et al. 2015). This result indicates a lower negative selection on the protein sequences compared to the rest of the genome. The accumulation of these variants with potential impact on gene functions might explain the high number of fitness and expression QTL detected in this region.

### High density of local regulatory variants detected is related to high genetic diversity, especially in the introgressed region

As high number of eQTL of the chromosome C hotspot affect expression of genes within the introgressed region, we wanted to investigate if the density of these “local”-effects is higher than in the rest of the genome. Hence we considered eQTL located at proximity of the disregulated gene to be “local” in the rest of the genome (see method). While these local-eQTL can be due to a mutation that directly changes the gene expression in *cis*, for example by changing a binding site in the promoter, they can also be explained by mutation located in a nearby gene acting in *trans*, or mutation disrupting the function of the affected gene, thus changing its expression through a retro-control (Ronald et al. 2005). About 54% of the eQTL detected correspond to local-eQTL (1,188). As observed in other eQTL analysis, local-eQTL generally have a stronger effect than non-local eQTL (999 distant-eQTL) (table S5 and figure S7B).

As expected, the density of local-eQTL is much higher in the introgressed region (160 per Mb) compared to the rest of the genome (100 per Mb). We proposed that the high genetic diversity of the region is the main factor explaining this higher density, increasing the chance of variants affecting expression in *cis*. We confirmed this observation by comparing the density of local-eQTL across different crosses from multiple yeast species with various genetic diversity. We used eQTL data from a cross between RM11.1a and BY4741 *S. cerevisiae* strains, as well as between 968 and Y0036 *S. pombe* strains, with a genetic diversity of 0.5% and 0.05%, respectively (Clément-Ziza et al. 2014; Albert et al. 2018). Interestingly, we observed a strong correlation between the density of genetic variants and local-eQTL (Figure 3, Pearson R^2^: 0.98, p-value: 0.001). This correlation allows us to explain the high density of local-eQTL in *L. kluyveri* and especially in the introgressed region. However, other factors are expected to impact the strong relationship observed here, such as the genome density and complexity of local regulation. Moreover, this correlation can be made only using dataset with similar detection power, as an increased power allows to observe very small genetic effects. Consequently, local-eQTL have been detected for 50% of *S. cerevisiae* genes using a population of 1,012 segregants (Albert et al. 2018).

**Figure 3.**
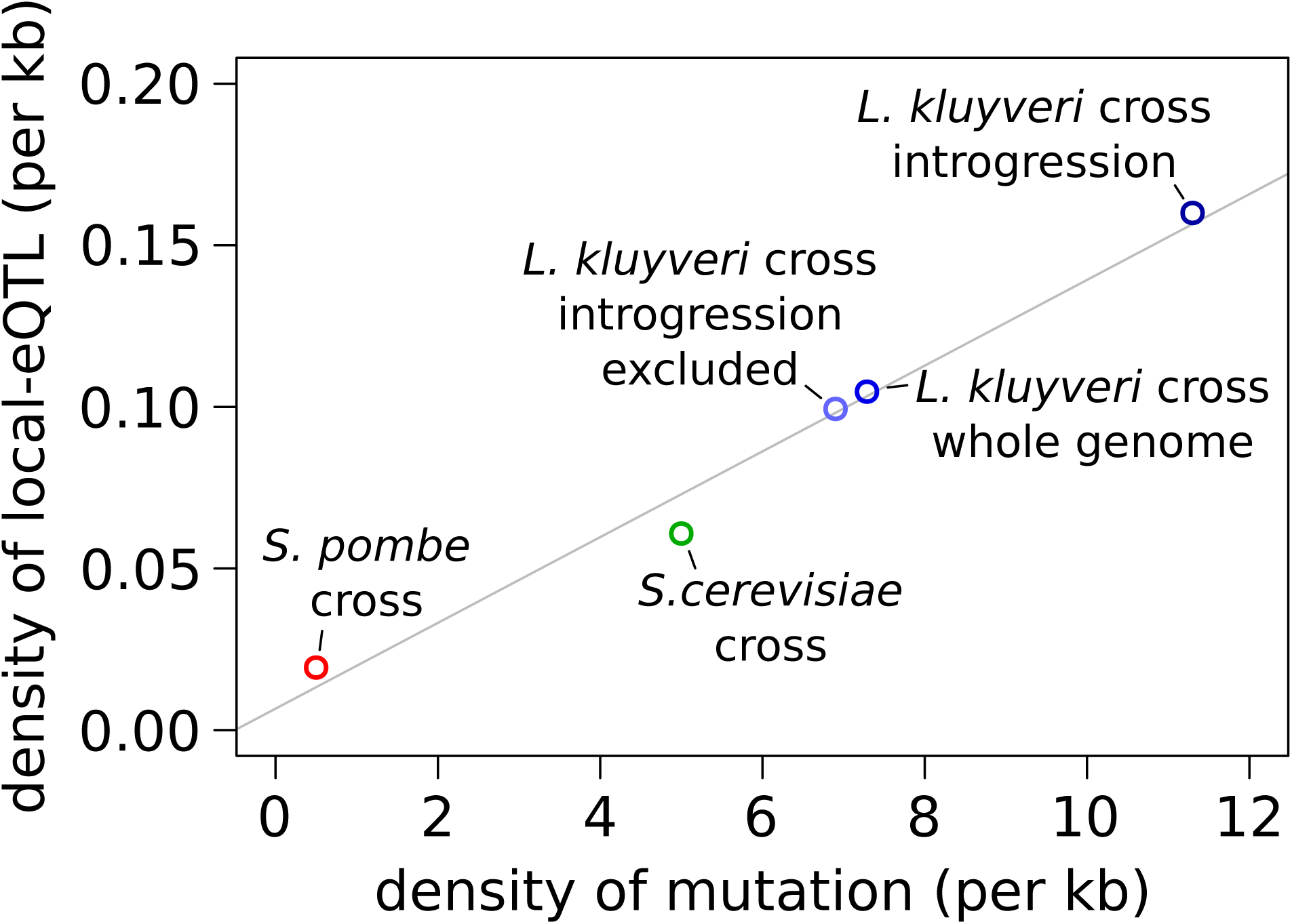
Comparison of the relationship between the density of mutation and the density of local eQTL across three yeast species. The grey line corresponds to a linear regression model (Pearson R ^2^ = 0.98, p-value = 0.001). Only the 2,000 strongest eQTL were considered among the 36,498 eQTL detected in the *S. cerevisiae* cross (see method). The data from the current study is indicated in shades of blue (*L. kluyveri*) compared to other studies in green (*S. cerevisiae*, (Albert et al. 2018)) and red (*S. pombe*, (Clément-Ziza et al. 2014)).

### Loci with major regulatory effect such as the one detected on chromosome E appear to be a trend across yeast species

The major QTL hostpot located on the chromosome E, containing 23 fitness QTL and about half of the detected distant-eQTL (496), influences resistance to various environmental stress. Consequently, we suspected that a part of the affected genes will be involved in the general environmental stress response (ESR) (Gasch et al. 2000; Brion et al. 2016). Indeed, we found a significant enrichment of 101 ESR genes among the 516 affected genes (Fisher’s exact test p-value: 7.6e^-4^). The impacted genes are also functionally enriched for electron transport and membrane-associated energy conservation (FunSpec p-value: <1e^-14^), tricarboxylic-acid pathway (FunSpec p-value: 7.1e^-7^), transcription (FunSpec p-value: 3.2e^-6^), and nucleus transport (FunSpec p-value: 1.6e^-4^). However, based on the limited annotation of *L. kluyveri*, no clear genes or variants can be proposed as directly responsible for this QTL hotspot (Figure S9).

The striking feature of this hotspot is the high number of gene expressions impacted by it (more than 500). However, similar hotspots have been found in other species such as in *S. cerevisiae* and *S. pombe* eQTL studies, where a major regulatory hotspot can be observed and narrowed down to the *MKT1* (D30G variant affecting 460 genes) and *SWC5* (Frame-shift affecting 610 genes) genes, respectively. However, these genes are involved in non-overlapping pathways (Figure S10). *MKT1* is involved in mRNA regulation and mitochondrial stability (Wickner 1987; Smith and Kruglyak 2008), and causes change of mitochondrial ribosomal protein and other mitochondrial proteins. By contrast, *SWC5* (component of the SWR1 complex) affects the expression of genes involved in cell cycle and chromosomal modification.

Finally, while many genes are affected by the hotspot located on the chromosome E, it tends to have a marginal effect on their expression variance (on average 39% of the variance explained) and on the growth capacity in the environment used for RNA extraction (7.5% of the growth variance explained in rich medium). Similarly in *S. cerevisiae, MKT1*-D30G only explains 19% of the variance of growth on minimum medium, and on average 23% of the variance of gene expression (Albert et al. 2018).

Overall, major regulatory hotspots seems to be a trend conserved across yeast species. While they are generally associated to a growth defect, the set of genes affected are specific to each case. Changing the environmental conditions likely creates a different major hotspot due to another genetic variant with a critical role catering to this new condition (Smith and Kruglyak 2008).

## Discussion

Dissecting the genotype-phenotype relationship remains a major challenge. Substantial improvement and availability of whole genome sequencing of large populations allows a more systematic analysis of the genetic origins of phenotypic diversity. However, large genomic properties such as introgression events are still difficult to be exhaustively identified and consequently they are usually overlooked even if they are known to have an impact on the phenotypic landscape.

In this context, we sought to assess the phenotypic effects of a 1-Mb introgressed region present in a *L. kluyveri* population. Interestingly, this region has a significant impact on the phenotypic landscape as illustrated by the fact that it corresponds to a QTL hotspot, composed of 23 fitness QTL and 247 eQTL. In addition, the absence of recombination in this introgressed region that determines mating-types genetically links all the genes and, consequently, all phenotypic characteristics their alleles might induce are linked to the *MAT* locus. As a direct result, *MAT*a and *MAT*α strains in the mapping population have very distinct phenotypic profiles and can perfectly be separated on a principal component analysis. This observation clearly highlights the presence of a sexual dimorphism in *L. kluyveri, i.e.* the association between the sex locus and some traits, which are initially unrelated to the determination of the sex.

In many plants and animals, the sex determination system is carried by chromosomes of different structure that does not recombine (X/Y or Z/W chromosomes). Among other fungi, only few cases of recombination suppression around the *MAT* locus have been observed. In the filamentous fungi *Neurospora tetrasperma*, a 7-Mb non-recombining regions, with over 1,500 genes, containing the *MAT* locus have been linked to a progressive accumulation of inversions between the two mating types (Menkis et al. 2008; Samils et al. 2013; Idnurm et al. 2015). This region also contained the centromere of the chromosome and ensure segregation of the mating types at the first meiotic division. Similarly, in the genome of *Podospora anserine*, a region of 837 kb (229 genes) around the *MAT* locus is depleted of recombination. In this species however, the region from both mating types remained colinear, and other yet-to-defined factors are responsible of the recombination depletion (Grognet et al. 2014). In the *Cryptococcus* and *Microbotryum* spp., absence of recombination allow genetic linking of two regions carrying pheromome-receptor genes and homeodomain genes, conserving a 50% compatibility in the progeny (Hsueh et al. 2006; Fontanillas et al. 2015; Idnurm et al. 2015).

With the generation of a mapping population in *L. kluyveri* for QTL identification, we simultaneously discovered a large region unable to go through recombination and observed a striking consequence on a large panel of phenotypes, leading to sexual dimorphism. Those phenotypes are poorly related, involving resistance to environmental stress (*e.g.* H_2_O_2_ or CaCl_2_) or growth in nutrient limitation (nitrogen limited media). Our eQTL results also revealed that the expression of many genes are affected by variants linked to the *MAT* locus, and can be considered as *MAT*-controlled genes. However, the majority of them have no direct link with pheromone production, sensing and mating. While the key element leading to this dimorphism is the absence of recombination on this 1-Mb region, it is still unclear what are the molecular origin of it. In any case, this region has been acquired through an introgression event, likely via the *E. gossypii* species (Landerer et al. 2019), and our results represent an unforeseen consequences of such event in the dynamics of evolution of species.

Importantly, such genetic load in the *MAT* region with no recombination was similarly observed in the other fungus species with the same properties (Samils et al. 2013; Grognet et al. 2014; Fontanillas et al. 2015). For almost two decades, it has been proposed that absence of recombination leads to genetic degeneration of the region around the *MAT* locus (Charlesworth and Charlesworth 2000). Indeed, all variants in these regions are genetically linked, preventing independent selection and elimination of deleterious mutations. The same phenomenon can be proposed in *L. kluyveri* were *de novo* mutations in the introgressed region would hijack the conservation of the *MAT* allele required for sexual cycle. This hypothesis can explain the high genetic diversity in the sequenced population, and the higher dN/ dS values for the genes located in this region (Brion et al. 2015; Friedrich et al. 2015). To confirm this hypothesis, we aim, in the future, to evaluate the rate of novel mutation along the genome in this species, through mutation accumulation experiments (Lynch et al. 2008).

Overall, by demonstrating that variations for an extensive set of phenotypes are linked to the *MAT* locus, we have shown the importance of given introgression features, *i.e.* a higher diversity and recombination suppression, in shaping the phenotypic diversity of the species. In addition, we also highlighted, for the first time, the possibility of sexual dimorphism in a budding yeast.

## Materials and methods

### Strains construction and F1 mapping population

The construction of the mapping population was previously described in (Brion et al. 2017). Briefly, the parental strains NBRC10955a *MAT*a chs3Δ and 67-588 *MAT*α *chs3Δ* were crossed on standard media (YPD: yeast extract 1%, peptone 2%, glucose 2%, agar 2%) and a hybrid was isolated. The deletion of *CHS3* allows for a better tetrad dissection (Sigwalt et al. 2016). The hybrid strain was put on sporulation media (potassium acetate 1%, agar 2%) for about a week. After digestion of the asci using zymolyase (0.5 mg/mL MP Biomedicals MT ImmunO 20T), 120 tetrads were dissected using a MSM 400 dissection microscope (Singer instrument). From the 120 tetrads dissected, 57 were completely viable. The 198 strains from 49 full-viable tetrads and two from a 50% viable tetrad were sequenced. The genomic DNA extracted using MasterPure Yeast DNA Purification Kit (tebu-bio: now Lucigen) was sequenced using Illumina HiSeq 2000 technology with 100 bp paired-end libraries. The reads were aligned to the CBS3082 reference genome using BWA (-n 8 -o 2 option) and the average coverage was around 70x for all the segregants. We used the same genetic markers as in our previous study. These 58,256 markers (SNPs) were selected to be highly reliable (see criteria in (Brion et al. 2017)). An allelic origin was assigned for each marker position of each segregant using roughly the same criteria of reliability as for the genetic markers determination. Raw data are available on the European Nucleotide Archive (http://www.ebi.ac.uk/ena) under accession number PRJEB13706.

### Growth condition and phenotyping

From the 198 sequenced segregants, 182 were allocated randomly on two 96 well plates, along with the parental strains, NBRC10955a and 67-588, and the hybrid NBRC10955a x 67-588. These two 96 well plates were used twice as matrix to create a YPD agar plate with 376 colonies (duplicates of the 182 segregant and four replicates for each parent on a 384 format). This plate was used as pregrowth for the fitness assessment. Cells for the colonies were transferred to two agar plates containing the test media using an automated pinning robot (Singer Instrument ROTOR HDA). This allows for a total of four replicates per strain and per condition. The list of the 64 growth conditions used are described in table S1: 2 different temperatures, 8 different carbon sources, 12 different unique nitrogen sources, 20 toxic compounds, including solvent, ions, or antifungal, at different concentrations, and 4 different pH. The pH was adjusted using a neutral citrate buffer. The base media was YP (yeast extract 1%, peptone 2%, agar 2%), YNB (yeast nitrogen base without nitrogen 1.7 g.l^-1^, glucose 2%, agar 2%), or SC (yeast nitrogen base with nitrogen 6.7 g.l^-1^, SC complement with all amino-acid 2 g.l^-1^, glucose 2%, agar 2%). The plates were incubated at 30°C, unless indicated otherwise in table S1 for 40 hours. An image of the scan of the plate was analyzed on R-gitter to obtain the area of each colony (data in table S2). The quantitative fitness value was computed as the ratio between the tested media and the reference media, with the exception of YP glucose 2% where the raw colony size was used for quantitative value.

### Generation of F2 mapping population

Using the genotyping data, we determined the mating type of the F1 segregant strains. We generated 69 crosses between *MAT*a and *MAT*α strains. The paired strains were chosen in order to keep an even ratio of both parental alleles. The set of hybrids were put on sporulation media (potassium acetate 1%, agar 2%) for three to seven days at 30°C. The asci of the tetrads were digested for 30 min using zymolyase (0.5 mg/mL MP Biomedicals MT ImmunO 20T) and about 10 tetrads per cross were dissected using a MSM 400 dissection microscope (Singer Instrument). From 50 full viable tetrads, we selected 50 spores (one per tetrad) to constitute the F2 population. We controlled for the 50 F2 strains to be haploid using propidium iodine DNA staining and flow cytometry.

### RNA extraction and sequencing

The transcriptome profiling was performed during exponential growth in rich media. The 50 F2 segregants and the parental strains were pregrown in YPD overnight. A flask of 30 mL of YPD was inoculating at a final optical density (OD600 λ = 600nm) of 0.05. The OD600 was monitored and, when it reached 0.350 to 0.450, 7 mL of the culture were filtered, washed with sterile water, and immediately frozen in liquid nitrogen and store at −80°C.

For RNA extraction, the cells were lysed in lysis buffer (EDTA 10mM, SDS 0.5%, TricHCl 10mM pH7.5) and aqua-phenol (MP AQUAPHEO01) at 65°C for one hour. The aqueous phase was purified from protein trace by a centrifugation in PLG tube (5prime 2302830). After precipitation in 70% ethanol overnight, the RNA was purified using the RNeasy kit (Qiagen 74104, Venlo, Limburg, Netherlands), and cleaned with DNase treatment (Cat No. 18068-015, Invitrogen, Carlsbad, CA, USA). The quality was controlled by a migration on agarose gel and RNA was quantified using spectrophotometry (NanoDrop ND-1000).

The sequencing of the 52 RNA samples was performed at the Gene Core Facility - EMBL (Heidelberg Germany) using multiplexed Illumina HiSeq2000 sequencing (50bp non-oriented single end reads). Raw reads data are available on the European Nucleotide Archive (http://www.ebi.ac.uk/ena) under accession number PRJEB32833. The normalization was performed as described in (Brion et al., 2015). Briefly, after excluding low expressed genes (number of reads lower than 32 for all samples), inter-sample normalization was performed using R and DEseq2 (R Core Team 2013; Love et al. 2014). Gene expressions from the retrogressed region were normalized separately due to the bias caused by the higher GC content. The base 2 logarithm of the normalized reads count was used as quantitative value for the abundance of mRNA.

### Genotyping of the F2 population

Using RNA-seq data, the F2 population was genotyped by determining the allelic origin of 58,256 markers previously used for the F1 population. We mapped the reads from RNA-seq on the reference genome and identified variants using SAMtools (Li et al. 2009). An allelic origin was inferred at a marker position if (1) the coverage of the marker was more than 2X, (2) the associated allele frequency was 0 or 1, and (3) the base sequenced fit one of the two parental alleles. If not, a NA flag was assigned to the position. The marker was kept for the next step if (1) it was located in a coding region, (2) had less than 30% of the F2 population that display a NA flag on its position, and (3) showed an unbalanced allele frequency in the F2 population (more than 0.7 or less than 0.3). Finally, post filtration a total of 37,529 marker were retained. In order to reduce the computation time, we artificially reduced the number of markers to 3,779 using pseudo-marker every 3 kb. The allele origin assigned to the pseudo-markers was the predominant allele origin of all the markers within 4 kb of the pseudo-marker. Given the low recombination rate of *L. kluyveri*, this approximation did not generate a strong loss of accuracy.

### Fitness and expression heritability

The broad-sense heritability, *H*^*2*^, is defined as the part of the variance explained by heritable factors. In our case, it has been defined as the variance in the segregating population that is not explained by the experimental noise. For fitness data, the variance explained by the experimental noise was estimated using the mean of the variance across the parental replicates (four replicates for the two parental strains and the hybrid strain). For the expression data, we did not have replicates to estimate the noise, therefore we used the variance of expression across replicates from another dataset of expression across 24 *L. kluyveri* strains published in (Brion et al. 2015). This dataset corresponds to expression profiling in the same condition (mid-log phase, in liquid YPD) and follows exactly the same protocol for RNA extraction and sequencing.

### Fitness QTL and eQTL analysis

The linkage analysis between quantitative phenotype (growth ratio and mRNA abundance) and genotype was performed using R/qtl (model: normal, method: Haley-Knott regression) (Arends et al. 2010). The phenotypic QTL were obtained using 182 F1 segregants. The significant thresholds of LOD score (score of the logarithm of the odds) were defined individually for each phenotype performing 1000 permutations and setting the LOD score for which QTL were detected in only 50 permutations (FDR = 5%).

The expression QTL were also obtained using R/qtl (model: normal, method: Haley-Knott regression) based on the normalized read count of the 50 F2 segregants and the two parental strains. To define the false discovery rate, we performed 100 permutations of the complete dataset and processed the eQTL analysis. For a range of LOD score threshold, the FDR was estimated as the average number of detected false-eQTL across the 100 permutations divided by the number of eQTL detected in the true dataset (Figure S5). We used an overall significant LOD score threshold of 3.7 which allowed to detect 2,187 eQTL with an FDR of ∼ 4%. The use of a more stringent LOD score threshold of 4.4 would have led to a detection of 1,638 eQTL with an FDR of ∼ 1%. Any functional enrichment were tested using the online tool FunSpec (Robinson et al. 2002).

### Expression QTL from *S. cerevisiae* and *S. pombe* used for interspecies comparison

To compare the genetic control of gene expression across species, we used published data from eQTL analysis in *S. cerevisiae* and *S. pombe*. For *S. cerevisiae* eQTL data, we used the 36,498 eQTL obtained from the population of 1,012 segregants for the RM11.1a and BY4741 cross (0.5% diversity) (Albert et al. 2018). However, because the detection power is much higher due to the large population, we only focus on the best 2,000 eQTL (2,000 highest LOD score, LOD score > 39.3), estimating that a population of about 50 segregants would have allowed for the identification of these eQTL. For *S. pombe*, we used the 2,346 eQTL obtained from a cross between 968 and Y0036 (0.05% diversity) with 44 F2 segregants (Clément-Ziza et al. 2014).

## Acknowledgments

We are grateful to Gargi Dayama for insightful comments on the manuscript. We thank the BioImage platform (IBMP-CNRS, Strasbourg, France) for their support. This work was supported by the Agence Nationale de la Recherche (ANR-18-CE12-0013-02) and a European Research Council (ERC) Consolidator grant (772505). J.S. is a Fellow of the University of Strasbourg Institute for Advanced Study (USIAS) and a member of the Institut Universitaire de France.

## Supplementary Material

Supplementary_Figures.pdf

Supplementary_Tables.xlsx

