## Supplementary_Figures for "Pervasive phenotypic impact of a large non-recombining introgressed region in yeast"

Christian Brion, Claudia Caradec, David Pflieger, Anne Friedrich, and Joseph Schacherer

Université de Strasbourg, CNRS, GMGM UMR 7156, F-67000 Strasbourg, France

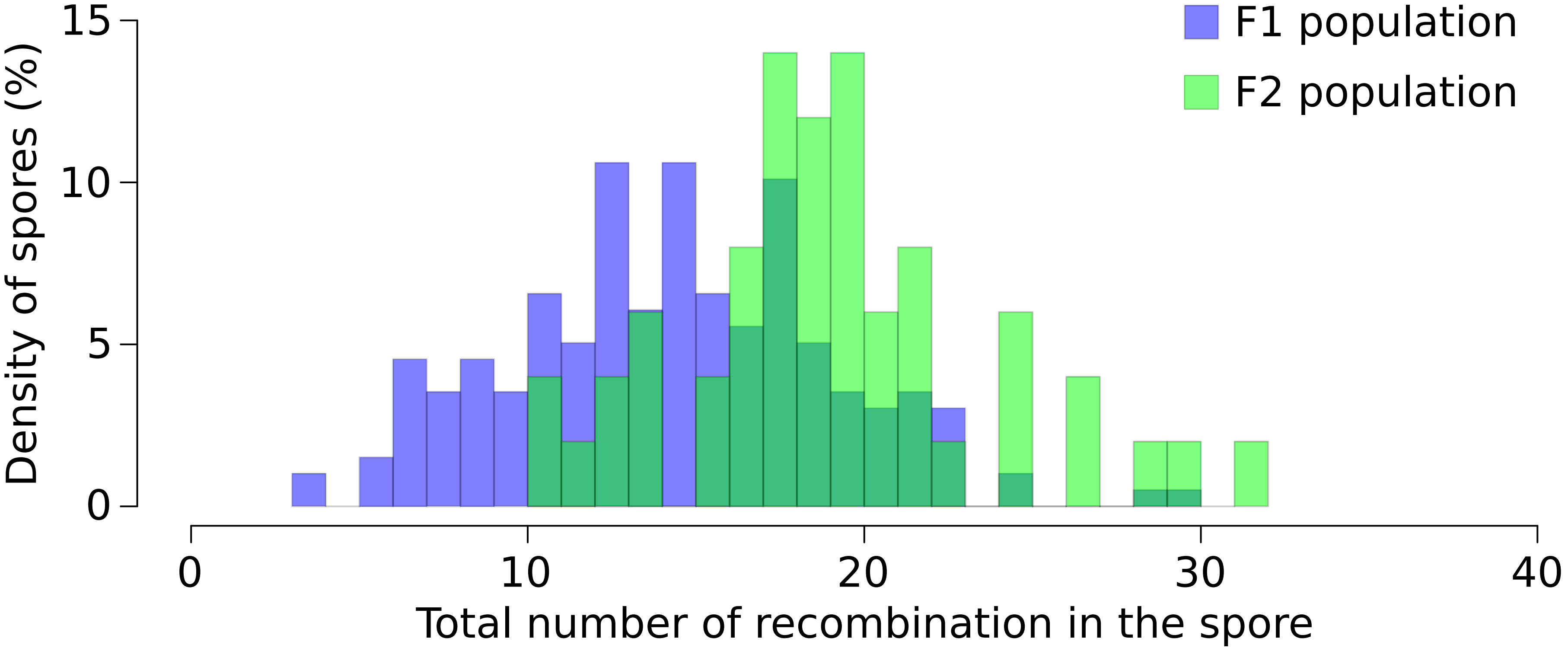


**Figure S1**. Distribution of the number of recombination per spore in the F1 and F2 populations.

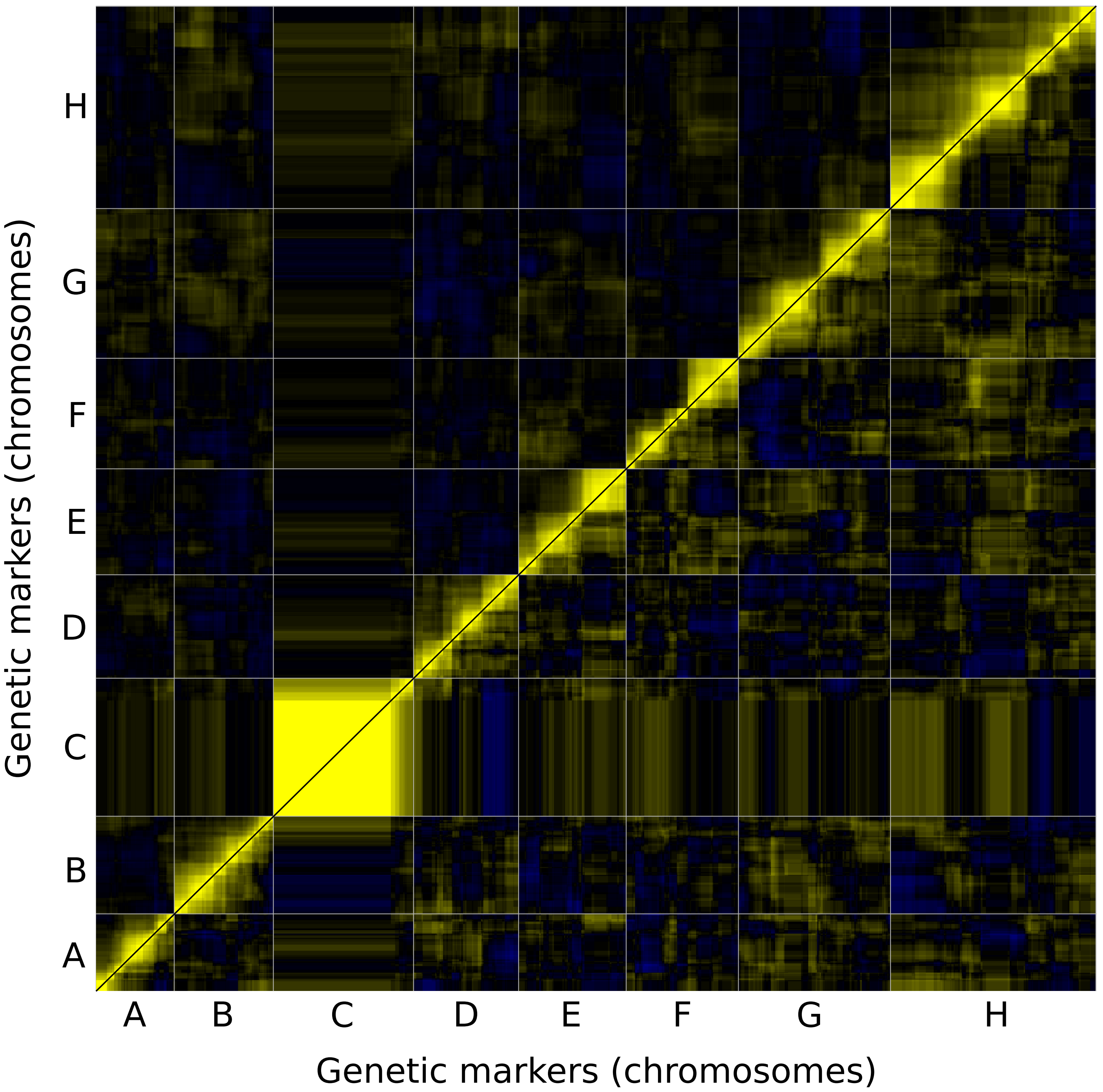


**Figure S2**. Linkage hit-map based on the pairwise correlation between markers in the F1 (up-left) and F2 (low-right) populations. Each pixel color represents the correlation coefficient between the allelic distribution across segregant of two markers. The low recombination rate generates block of markers, in yellow, with identical allelic distributions.

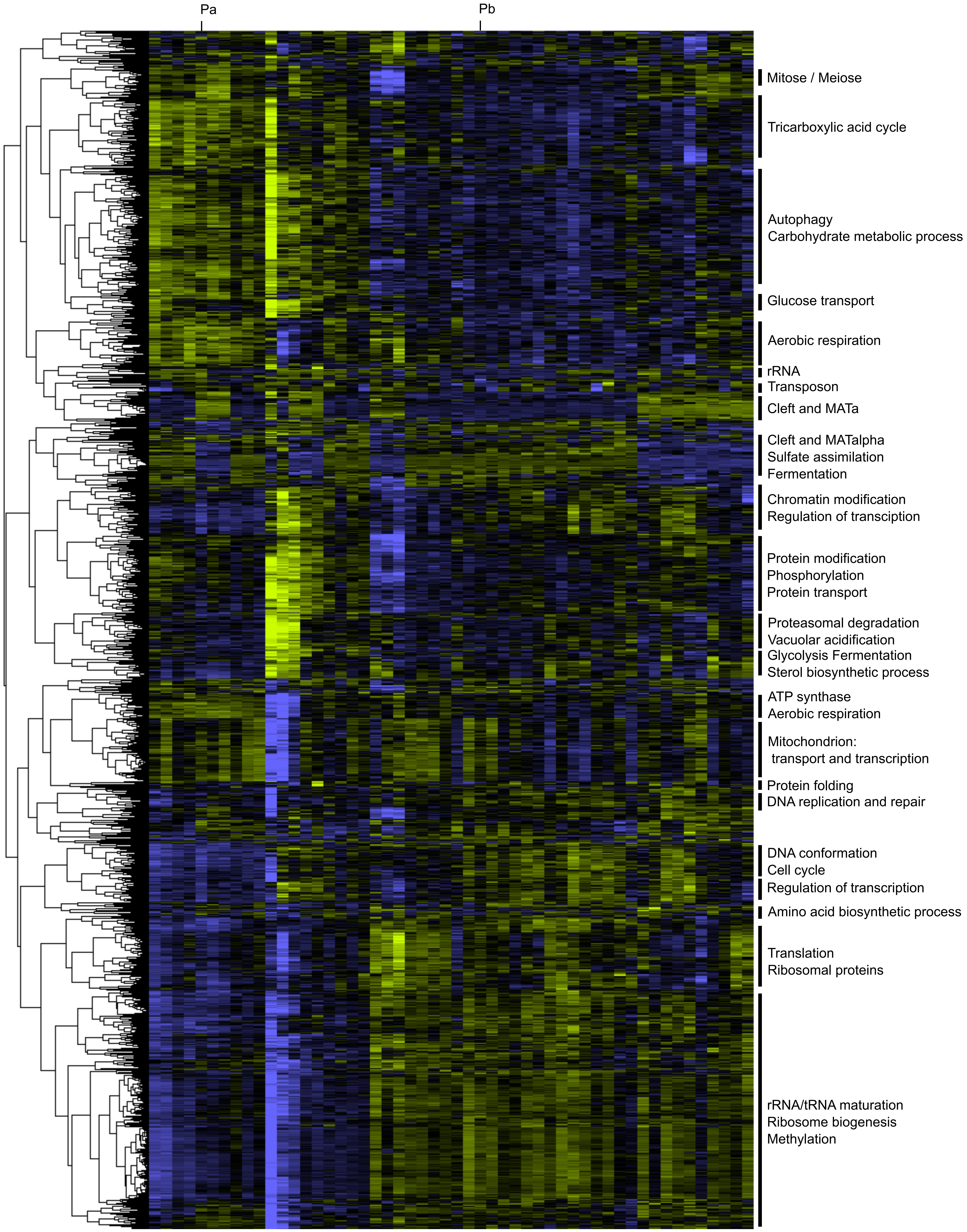


**Figure S3**. Clustering of expression profiles of gene across the 50 F2 segregants and the two parental strains, indicated as Pa for NBRC10955 and Pb for 87-588. Functional enrichment of the genes in each cluster are indicated on the right, based on Funspec (Robinson *et al.*, 2002).

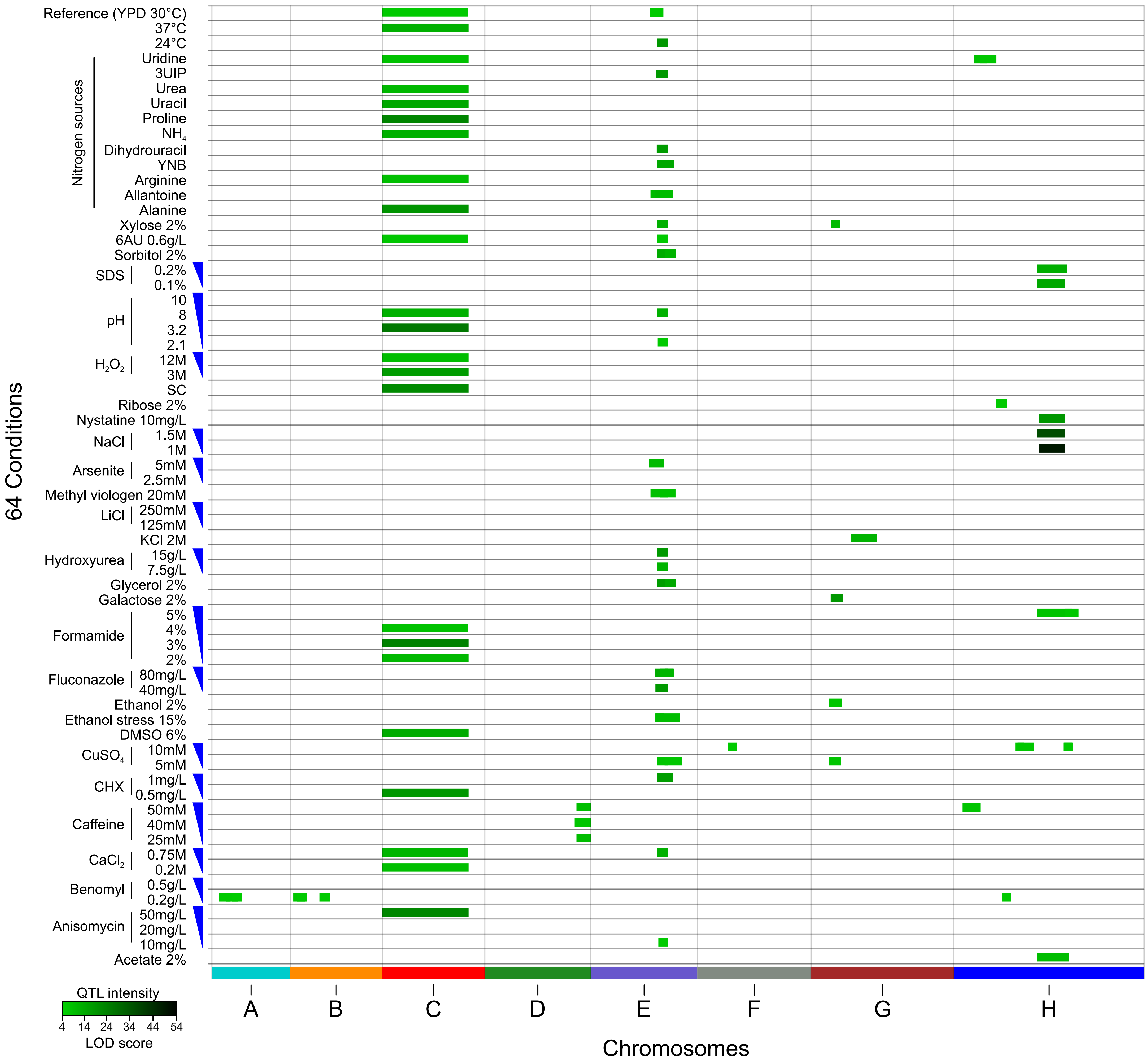


**Figure S4**. Position of the QTL detected across all the 64 conditions. The color intensity is correlated to the maximum LOD score.

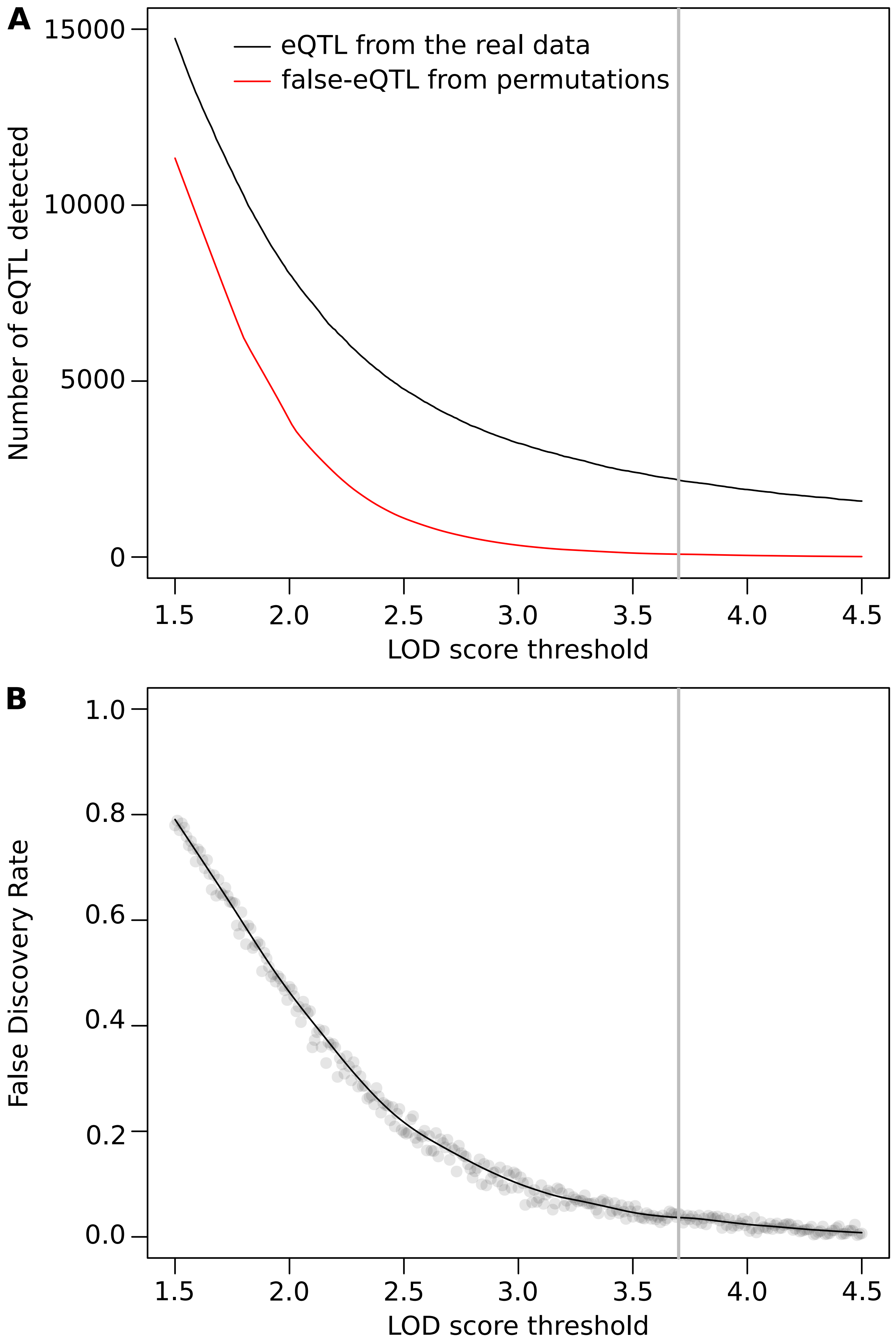


**Figure S5**. Defining eQTL false discovery rate using permutation data. A) Evolution of the number of eQTL detected in relation to the significant threshold of the LOD score. B) Decrease of the FDR due to the increase in the threshold of the LOD score. The FDR is defined as the average number of detected false-eQTL across the 100 permutations divided by the number of eQTL detected in the real dataset. The significant threshold of the LOD score used is indicated by the grey bar (3.7 LOD score) which corresponds to a FDR of 4%.

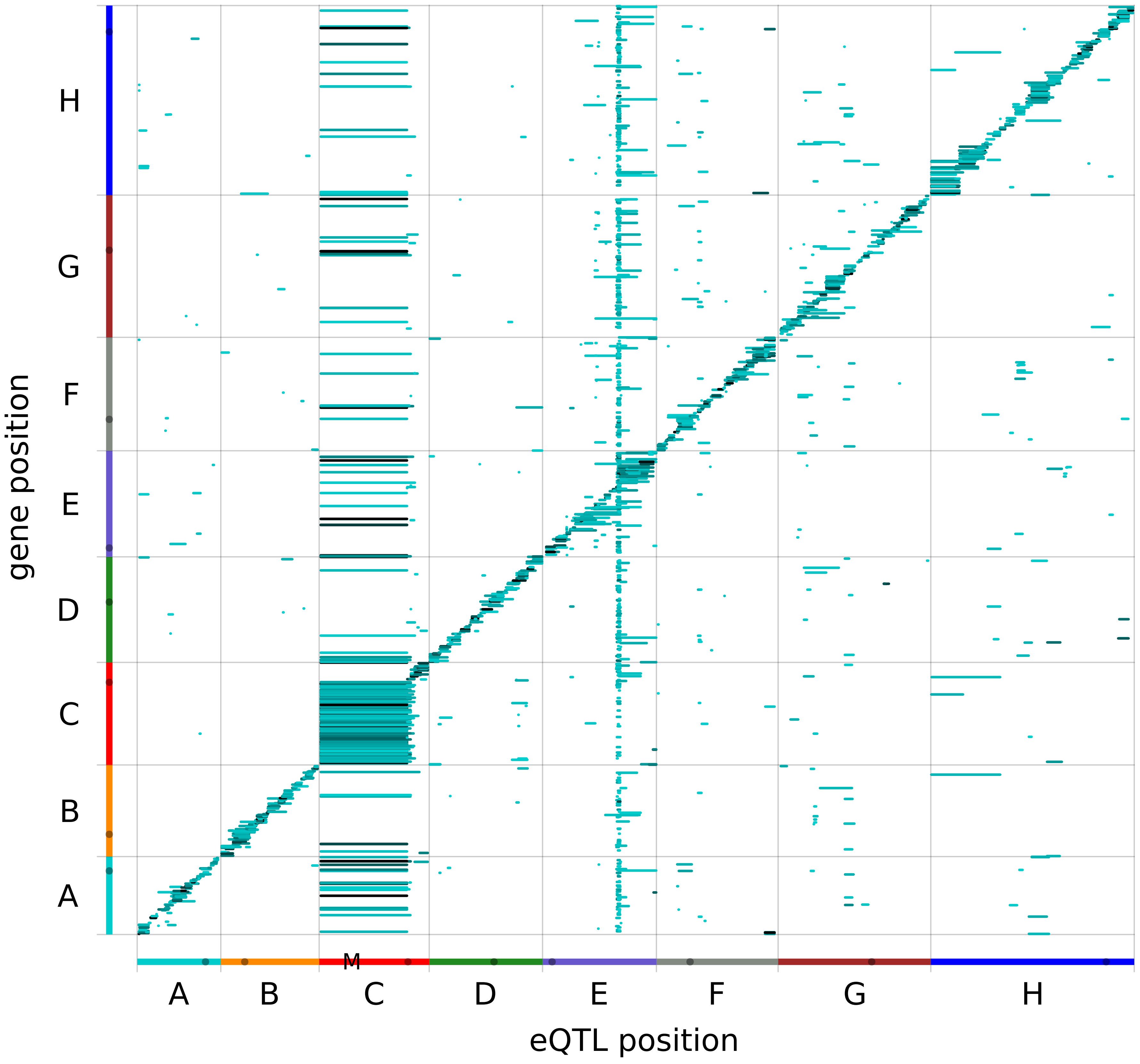


**Figure S6**. Relation between eQTL position (X-axis) and the location of the affected gene (Y-axis). Vertical overlap corresponds to hotspot of distant eQTL. The eQTL on the diagonal correspond to the local-eQTL. The color intensity is correlated to the maximum LOD score. The M indicates the position of the *MAT* locus.

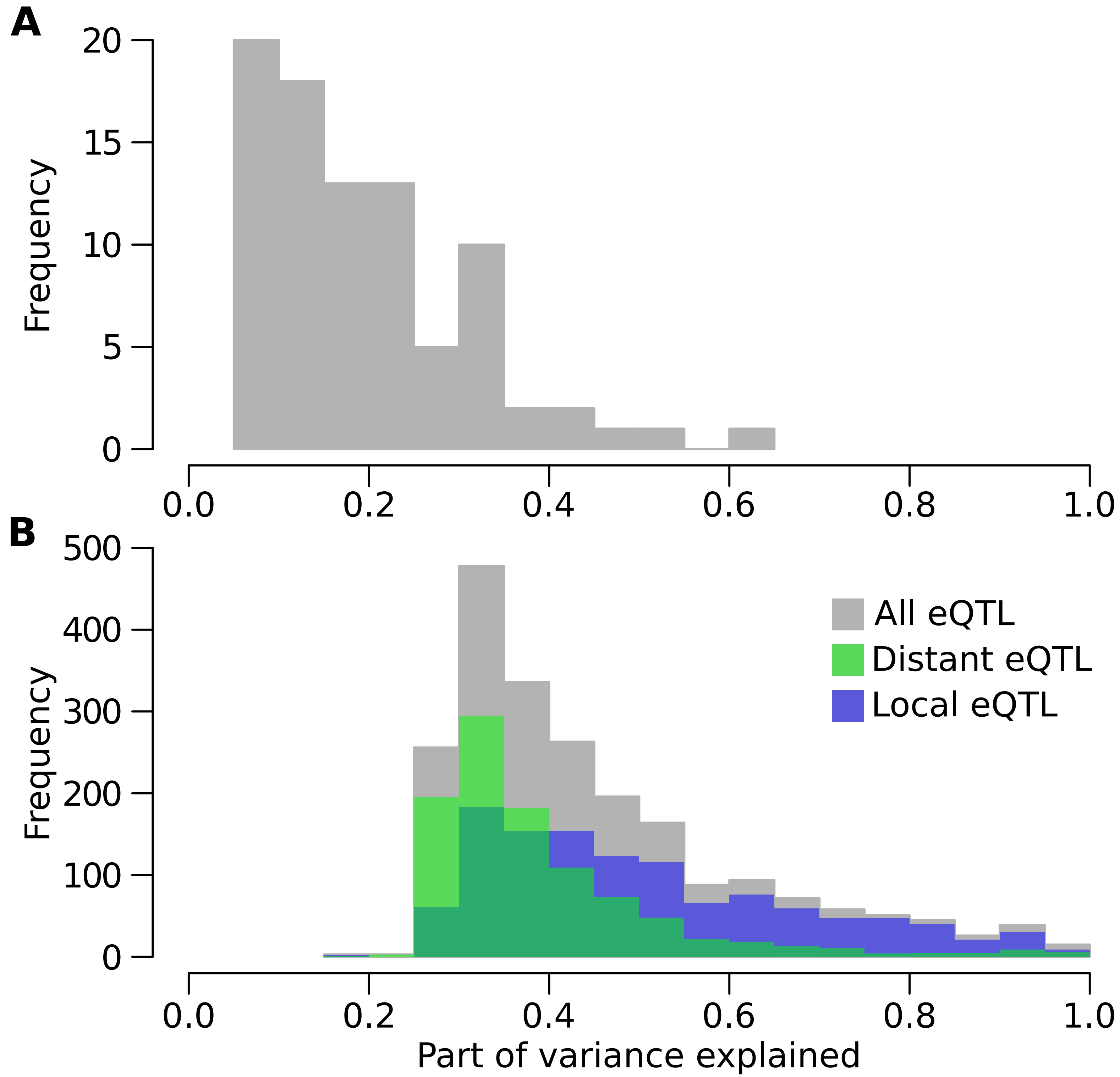


**Figure S7**. Distribution of the variance explained for each QTL (A) and eQTL (B) detected. The green and blue histogram correspond to distant and local eQTL, respectively.

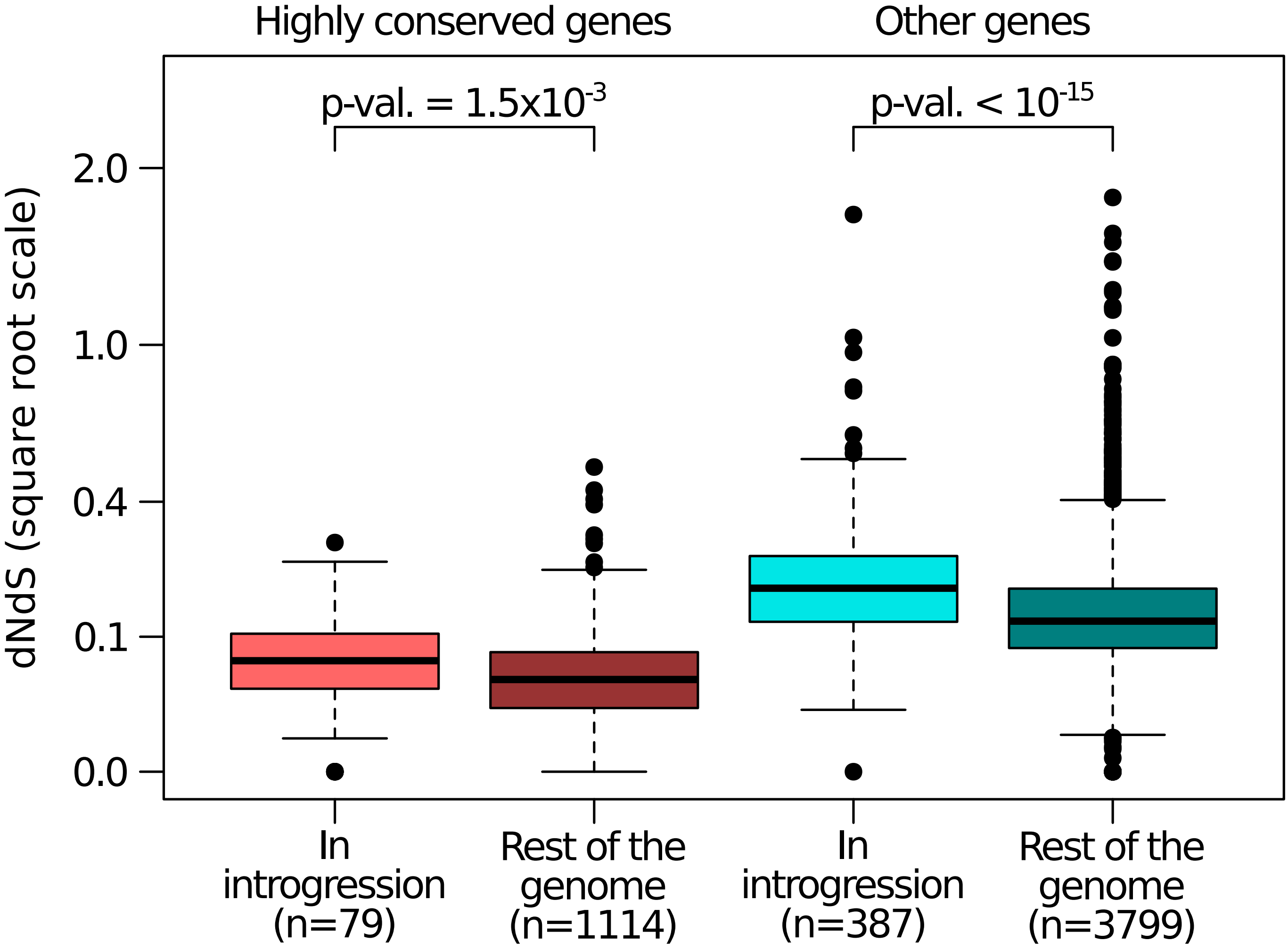


**Figure S8**. Significantly higher dNdS value for the genes in the introgressed region compared to the rest of the genome. Highly conserved genes are the *L. kluyveri* genes with more that 80% of protein similarity with their ortholog in *S. cerevisiae*. P-values correspond to a t-test.

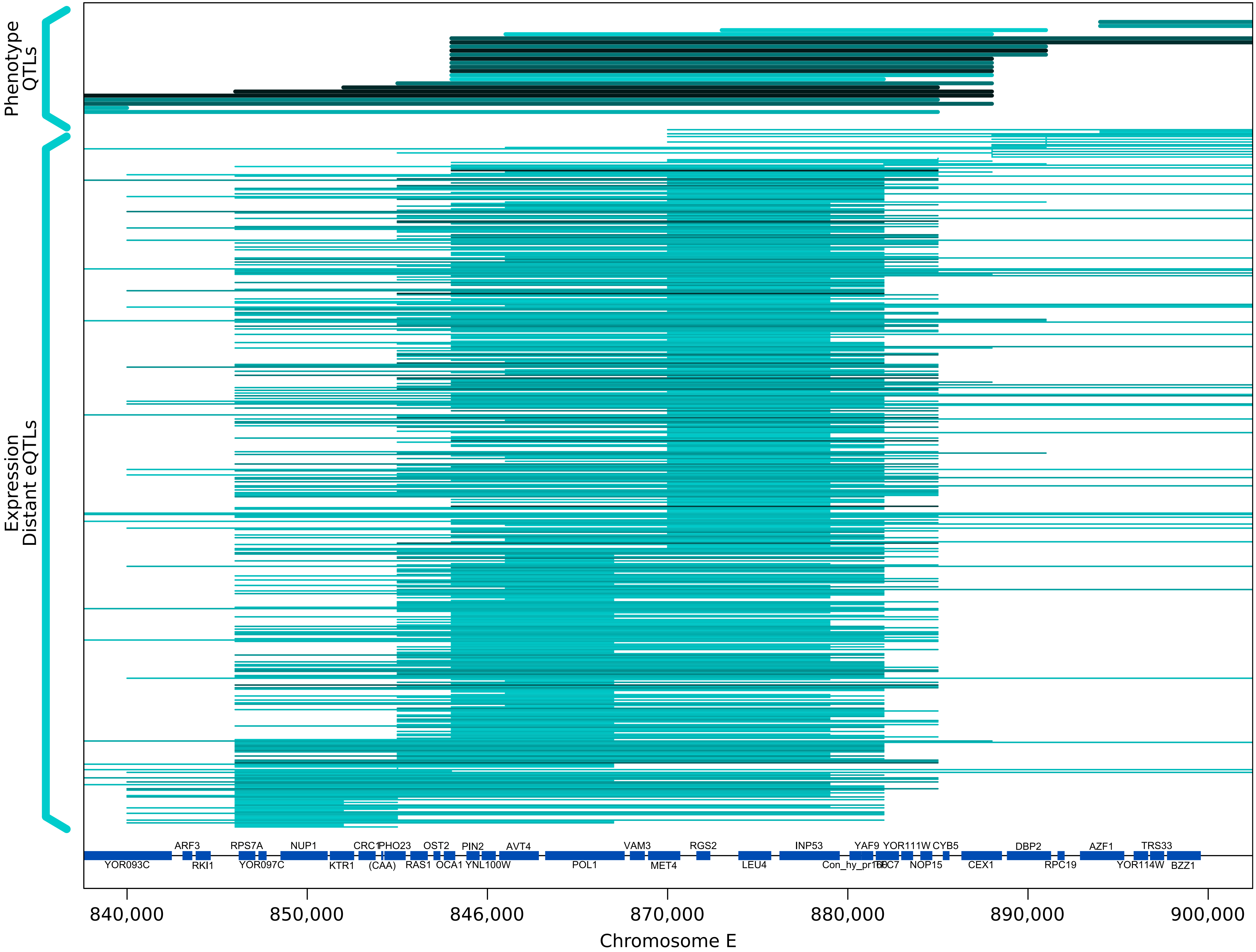


**Figure S9**. Position of the Phenotypic QTL (top) and eQTL (bottom) over the chrE:870k hotspot. The color intensity is correlated to the maximum LOD score.

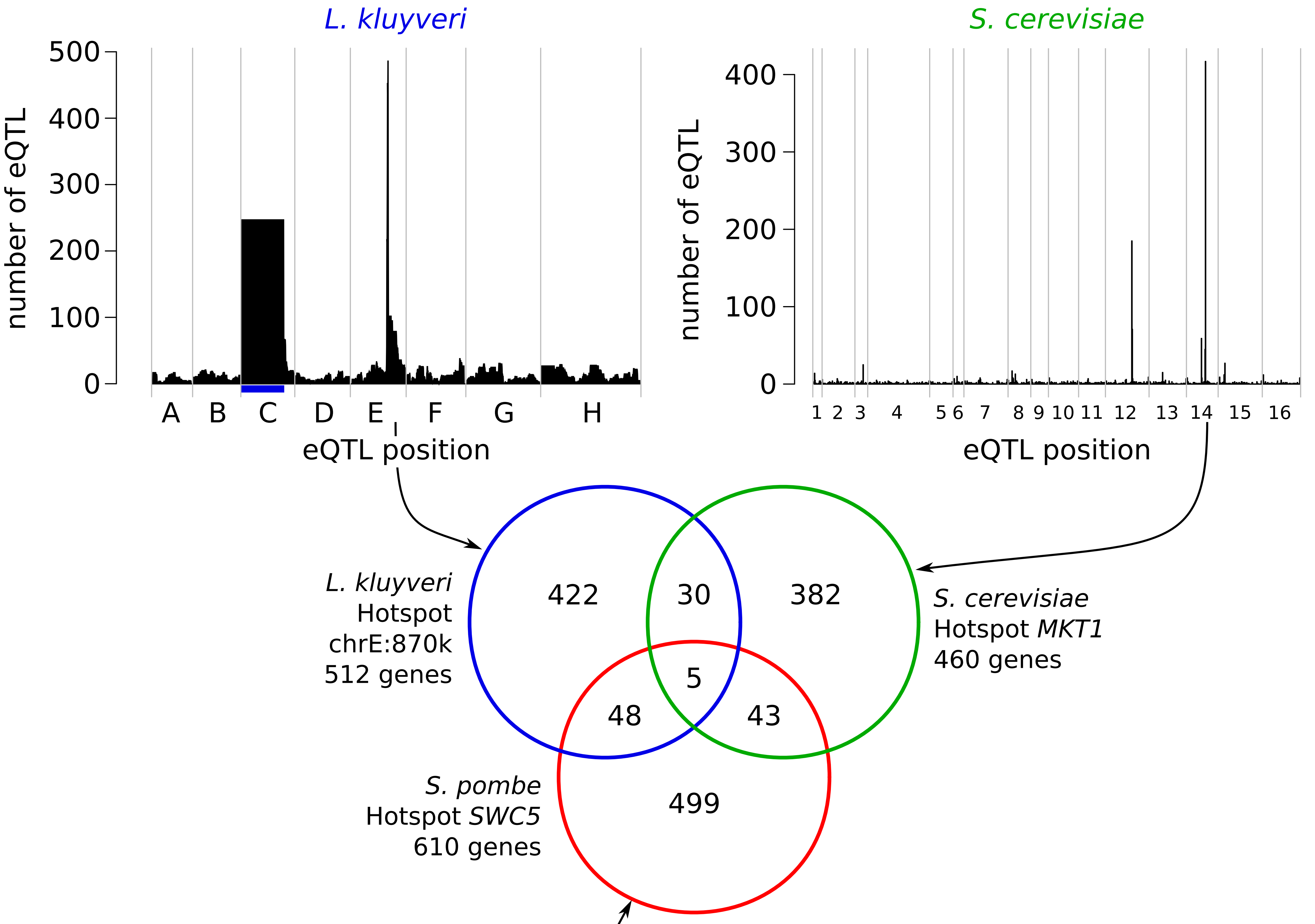


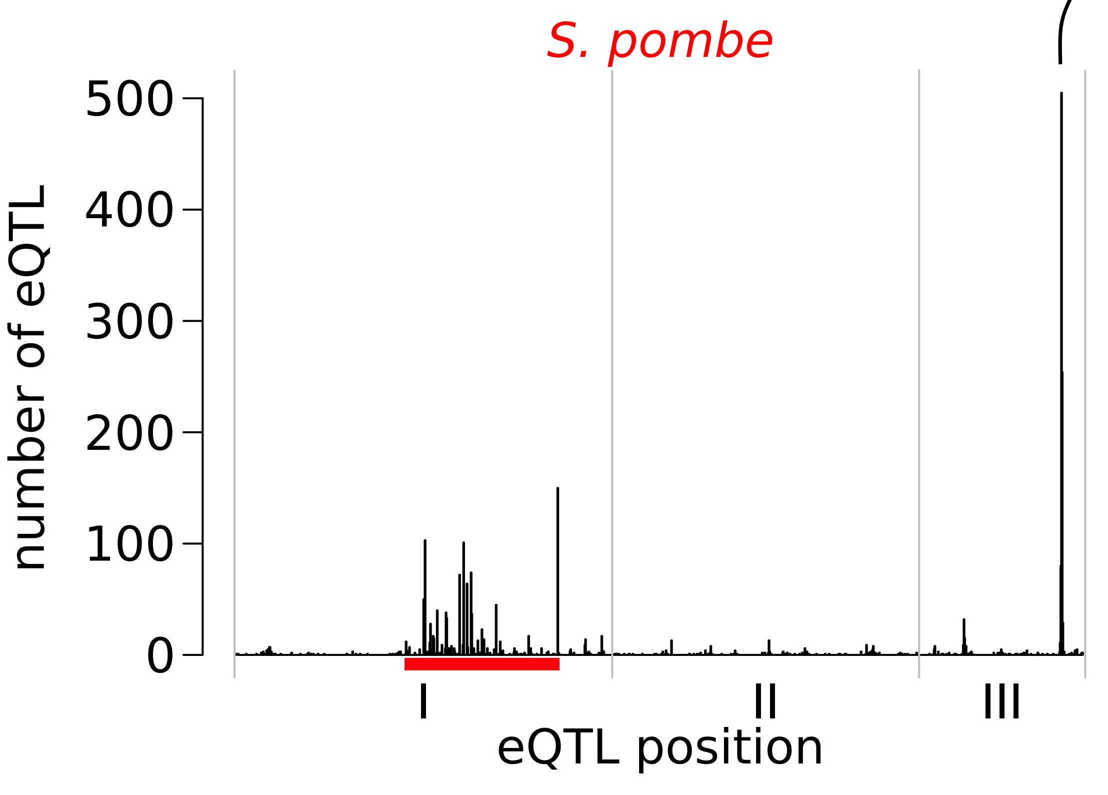


| study | major hotspot functional enrichment | p-value |
|---|---|---|
| *L. kluyveri* this study | electron transport | $<10^{-14}$ |
| | TCA | $7x10^{-7}$ |
| | transcription | $3.2x10^{-6}$ |
| *S. cerevisiae* Albert *et al.* | mitochondrial translation | $<10^{-14}$ |
| | aerobic respiration | $3.7x10^{-6}$ |
| *S. pombe* Clement-Ziza *et al.* | mRNA processing | $4.3x10^{-7}$ |
| | chromatin modification | $6.1x10^{-7}$ |
| | cell cycle | $10^{-5}$ |

**Figure S10**. Comparison of major regulatory hotspot across yeast species. The Venn diagram shows the absence of correlation between the genes affected by the major hotspot across the three species. The analysis is based on *S. cerevisiae* orthologous annotation for *L. kluyveri* and *S. pombe*. The functional enrichment is done with Funspec using for *L. kluyveri* and *S. pombe*, the *S. cerevisiae* orthologous annotation (Robinson *et al.*, 2002). On the density plots, the blue area for *L. kluyveri* indicate the introgressed region and the red area for *S. pombe* indicate a 2,230 kb inversion on chromosome I between the two parental strains. Only the 2,000 strongest eQTL were considered among the 36,498 eQTL detected in the *S. cerevisiae* cross (see method).
